# Homeostatic plasticity scales dendritic spine volumes and changes the threshold and specificity of Hebbian plasticity

**DOI:** 10.1101/308965

**Authors:** Anna F. Hobbiss, Yazmin Ramiro Cortés, Inbal Israely

**Affiliations:** Champalimaud Centre for the Unknown, 1400-038 Lisbon, Portugal.; Instituto de Fisiología Celular, Universidad Nacional Autónoma de México, Ciudad Universitaria, Circuito exterior s/n, Ciudad de México 04510, México.; Department of Pathology and Cell Biology in the Taub Institute for Research on Alzheimer’s Disease and the Aging Brain, Department of Neuroscience, College of Physicians & Surgeons, Columbia University, New York, New York, 10032, USA.

**Author notes:** **Corresponding author:** Inbal Israely. Electronic address.

## Abstract

Information is encoded within neural networks through synaptic weight changes. Synaptic learning rules involve a combination of rapid Hebbian plasticity with slower homeostatic synaptic plasticity (HSP) that regulates neuronal activity through global synaptic scaling. While Hebbian plasticity has been extensively investigated, much less is known about HSP. Here we investigate the structural and functional consequences of HSP at dendritic spines of mouse hippocampal neurons. We find that prolonged activity blockade induces spine growth, paralleling synaptic strength increases. Following activity blockade, glutamate uncaging-mediated long-term potentiation at single spines leads to size-dependent structural plasticity: smaller spines undergo robust growth, while larger spines remain unchanged. Moreover, we find that neighboring spines in the vicinity of the stimulated spine exhibit volume changes following HSP, indicating that plasticity has spread across a group of synapses. Overall, these findings demonstrate that Hebbian and homeostatic plasticity shape neural connectivity through coordinated structural plasticity of clustered inputs.

## Introduction

Hebbian synaptic plasticity, widely regarded as the leading biological mechanism for information storage, involves activity-dependent changes in synaptic connectivity (Malenka and Bear, 2004). Notably, these changes have a physical component and excitatory postsynaptic current is highly correlated with dendritic spine volume (Matsuzaki et al., 2001). However, left unchecked activity would lead to a positive feedback loop in which changes in synaptic weight are further reinforced by future events. This has been proposed to result in loss of functionality of the system, by either driving synapses towards saturation, or through silencing the population (Miller and MacKay, 1994). How neurons maintain stability in the face of destabilizing cellular and circuit events has been a longstanding question (LeMasson et al., 1993). One method neurons employ to resolve this is Homeostatic Synaptic Plasticity (HSP) (Turrigiano et al., 1998), a feedback mechanism through which a population of synapses can be maintained to function within optimal bounds. During HSP, a decrease in global activity drives a counteracting scalar increase in synaptic strengths to bring the network into an optimal functioning range, and conversely, network activity increases will promote synaptic weakening (Turrigiano, 2011). This scaling is instantiated in part by AMPA receptor trafficking to and from the post-synaptic density, as well as through presynaptic changes which modify neurotransmitter release, leading to concomitant changes in synaptic strength (Murthy et al., 2001; O’Brien et al., 1998). HSP has been observed both *in vitro* and *in vivo* (Desai et al., 2002), supporting a fundamental role for this mechanism in the proper functioning of a neural system.

It is well established that HSP modulates synaptic function; however, it is currently unknown how this process also correlates with spine structural changes. Moreover, how HSP affects the induction and longevity of subsequent forms of Hebbian plasticity at individual inputs remains to be determined. In this study, we examine how the induction of HSP affects the structure and function of hippocampal pyramidal neuron synapses, and what are the consequences of these changes for future Hebbian plasticity. We find that the induction of HSP through activity blockade leads to an overall increase in the size of spines, corresponding to a structural scaling that matches the functional scaling of synapses. Through precise 2-photon mediated glutamate uncaging, we further investigate how these homeostatic functional and structural plasticity changes impact the ability of individual inputs to undergo subsequent plasticity. We demonstrate that after HSP, spines express increased longevity of LTP, an increased growth rate after stimulation, and a reduced plasticity threshold. We find that HSP enhances the magnitude of synaptic potentiation through the preferential modulation of small spines and by promoting structural plasticity at clustered inputs following Hebbian activity at single synapses. Together, these changes provide a mechanism by which homeostatic plasticity can modulate synaptic efficacy and enhance future learning without compromising previously stored information.

## Results

### Structural correlates of homeostatic plasticity

Hebbian plasticity at an individual input is linearly correlated with volume changes in the corresponding spine (Govindarajan et al., 2011; Harvey and Svoboda, 2007; Matsuzaki et al., 2004). Since HSP is known to change the functional properties of the spine, we reasoned that homeostatic modifications of efficacy would be accompanied by structural plasticity of inputs. We induced HSP using prolonged activity blockade through Tetrodotoxin (TTX) inhibition of sodium channels, for either 0 h, 24 h, 48 h or 72 h (Figure 1a), as this has been shown to induce scaling of synaptic strengths in a variety of systems (Karmarkar and Buonomano, 2006; Turrigiano et al., 1998). We chose to conduct our experiments in mouse hippocampal organotypic slice cultures, which maintain a physiologically relevant tissue architecture and are amenable to genetic manipulation. To verify that functional synaptic scaling occurred, we recorded spontaneous miniature excitatory post-synaptic currents (mEPSCs) from both TTX treated and Control CA1 hippocampal pyramidal neurons at 48 h after the beginning of the HSP induction period (Figure 1b-d). All recordings were performed in the presence of acute TTX to block action potentials. As expected, we found a significant increase in mEPSC amplitude following 48 h of activity blockade (Figure 1b,c; Control = 18.6 ± 0.25 pA, TTX = 20.72 ± 0.28 pA, p = 1.55e-8 Mann Whitney). The distribution of the TTX mEPSCs scaled linearly to overlay with the control distribution (Figure 1d), in accordance with the synaptic scaling theory. We next investigated whether homeostatic plasticity leads to structural modification of synapses by live, 2-photon imaging of GFP-labelled CA1 dendrites (Figure 1e). The volumes of all visible spines within the image were measured using *SpineS*, a custom inhouse developed Matlab toolbox (Erdil et al., 2012). We found that spines from chronically TTX treated cells were significantly bigger than those from control cells beginning at 48 h (Figure 1e,f; Control = 0.136 ± 0.0062 μm^3^, TTX = 0.211 ± 0.0123 μm^3^). Surprisingly, as opposed to the linear functional scaling of mEPSCs seen in response to HSP, we found that structural scaling of spine volumes at 48 h are best fit to controls with a second order equation (Figure 1g). This superlinear scaling resulted from a preponderance of large spines after TTX treatment, for which correspondingly large mEPSCs were not observed.

**Figure 1.**
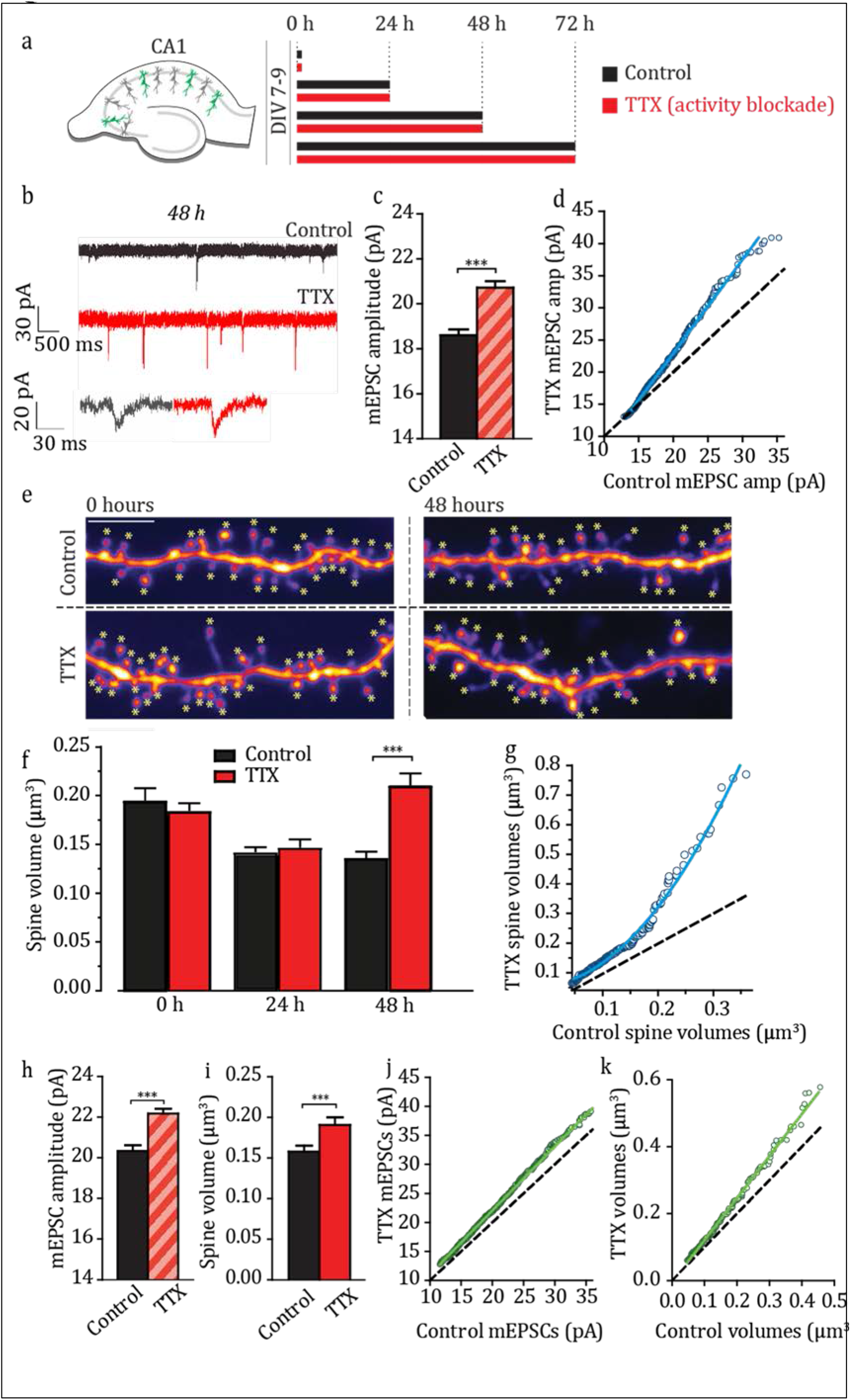
Spines undergo structural scaling following activity blockade. Experimental protocol to induce HSP in hippocampal organotypic slices. b) Example mEPSC recording traces from control and TTX-treated conditions at the 48 h time point. *Bottom:* representative individual mEPSCs. c) Quantification of mEPSC amplitudes after 48 h of activity blockade, represented as mean amplitudes ± SEM. ***p = 1.553e-08, Mann-Whitney. For all data presented, total number of spines or mEPSCs are represented by the first number (n) and followed by the number of independent neurons (N = in parentheses) from which they were collected. Number of mEPSCs (n) from independent neurons (N): Control = 552 (8), TTX = 742(8). d) Ranked control mEPSC amplitudes plotted against TTX condition at 48 h. Best fit line to the data: TTX = Control × 1.45 – 6.06. e) Representative images of dendrites at either 0 or 48 h after TTX treatment or in control media. Asterisks indicate all the spines that were analyzed. Scale bar = 5μm. f) Quantification of spine volumes for the indicated conditions. ***p = 0.91e-10, Two-way ANOVA, post-hoc Bonferroni. Number of spines (n) from independent neurons (N) for the 0 h, 24 h and 48 h timepoints: Control = 103(7), 245(8), 285 (7). TTX = 172 (7), 232 (5), 208 (6). g) Ranked control spine volumes plotted against TTX condition at 48 h. Best fit line to the data: TTX = Control^2^ × 5.44 + Control × 0.25 + 0.55. h) Quantification of mEPSC amplitudes after 72 h in TTX. ***p = 1.883e-15, Mann-Whitney. Number of mEPSCs (n) from independent neurons (N): Control = 1164 (10), TTX = 1302 (12). i) Quantification of spine volume after 72 h in TTX. ***p = 0.0008, Mann-Whitney. Number of spines (n) from independent neurons (N): Control = 258 (6), TTX = 214 (6). j) Ranked control 72 h mEPSC amplitudes plotted against TTX mEPSC amplitudes. Best fit line to the data: TTX = Control × 1.10 – 0.01. k) Ranked control 72 h spine volumes plotted against TTX 72 h spine volumes. Best fit line to the data: TTX = Control × 1.26 – 0.004).

After 72 h of activity blockade, synapses maintained the enhancement of mEPSC size (Figure 1h; Mean ± SEM: control 72 h = 20.35 ± 0.27 pA, TTX 72 h= 22.18 ± 0.239 pA, p = 1.88e-15, Mann-Whitney) and spine volume increases (Figure 1i; Mean ± SEM: control 72 h= 0.16 ± 0.0073 μm^3^, TTX = 0.19 ± 0.0093 μm^3^, p = 0.0008 Mann-Whitney). However, at this time, scaling was instantiated in a linear fashion at both the level of mEPSC amplitude and spine volumes (Figure 1j,k). This suggests that the expression of functional and structural scaling evolves dynamically over time in response to prolonged activity blockade.

### Reversibility of homeostatic plasticity mediated structural changes

Synaptic strength modifications that occur in response to activity blockade are reversible upon re-exposure of a circuit to activity (Desai et al., 2002; Wallace and Bear, 2004). We tested whether the structural changes that resulted following activity blockade were also reversible when activity is restored. After 48 h of homeostatic plasticity induction, by which time significant structural and functional scaling have occurred (Figure 1), we removed TTX allowing slices to resume activity and measured spontaneous firing using whole-cell patch clamp recordings (Figure 2a,b). Activity blockade for 48 h led to higher firing rates compared to untreated cells (Figure 2c; Control (median) = 1.33, IQ range = 0.11:18.83 spikes/min; TTX (median) = 13.33, IQ range = 11.5:21.06 spikes/min. p=0.03, Kruskal-Wallis post-hoc Dunn). To exclude the possibility that the higher firing rates we observed were due to a rebound after the acute withdrawal of TTX from the system, a set of control neurons were briefly incubated in TTX (2-4h, named “Acute TTX condition”). This manipulation did not significantly alter firing rates, which were similar to untreated controls (Figure 2c; Acute-TTX: med = 4.44, IQ range = 2.0:29.33 spikes/min vs Control no-TTX: Median = 1.33, IQ range = 0.11:18.83 spikes/min, p>0.9 Kruskal-Wallis post-hoc Dunn). Thus, the firing rate changes we observed were the result of HSP and did not arise from the acute withdrawal of TTX. We then investigated whether the circuit would return to its original level of activity once neurons were released from blockade. Indeed, after having removed TTX for 48 h, firing rates were indistinguishable from controls (Figure 2c; Control (median) = 1.33, IQ range = 0.11:15.56. TTX reversal (median) = 1.67, IQ range = 0.194:11.76. p>0.05, Kruskal-Wallace post-hoc Dunn). We then examined whether structural modifications accompanied these firing rate changes, and observed a similar reversal in spine volumes (Figure 2d,e). TTX treated spines significantly reduced in size 48 h after TTX removal (Figure 2e; mean volume ± SEM: TTX 48 h = 0.21 ± 0.0063 μm^3^; TTX 48 h reversal = 0.16 ± 0.0064 μm^3^; p=4.1852e-07). These findings support the idea that functional plasticity is matched by structural plasticity, and that spines sizes reversibly and accurately reflect the activity landscape.

**Figure 2.**
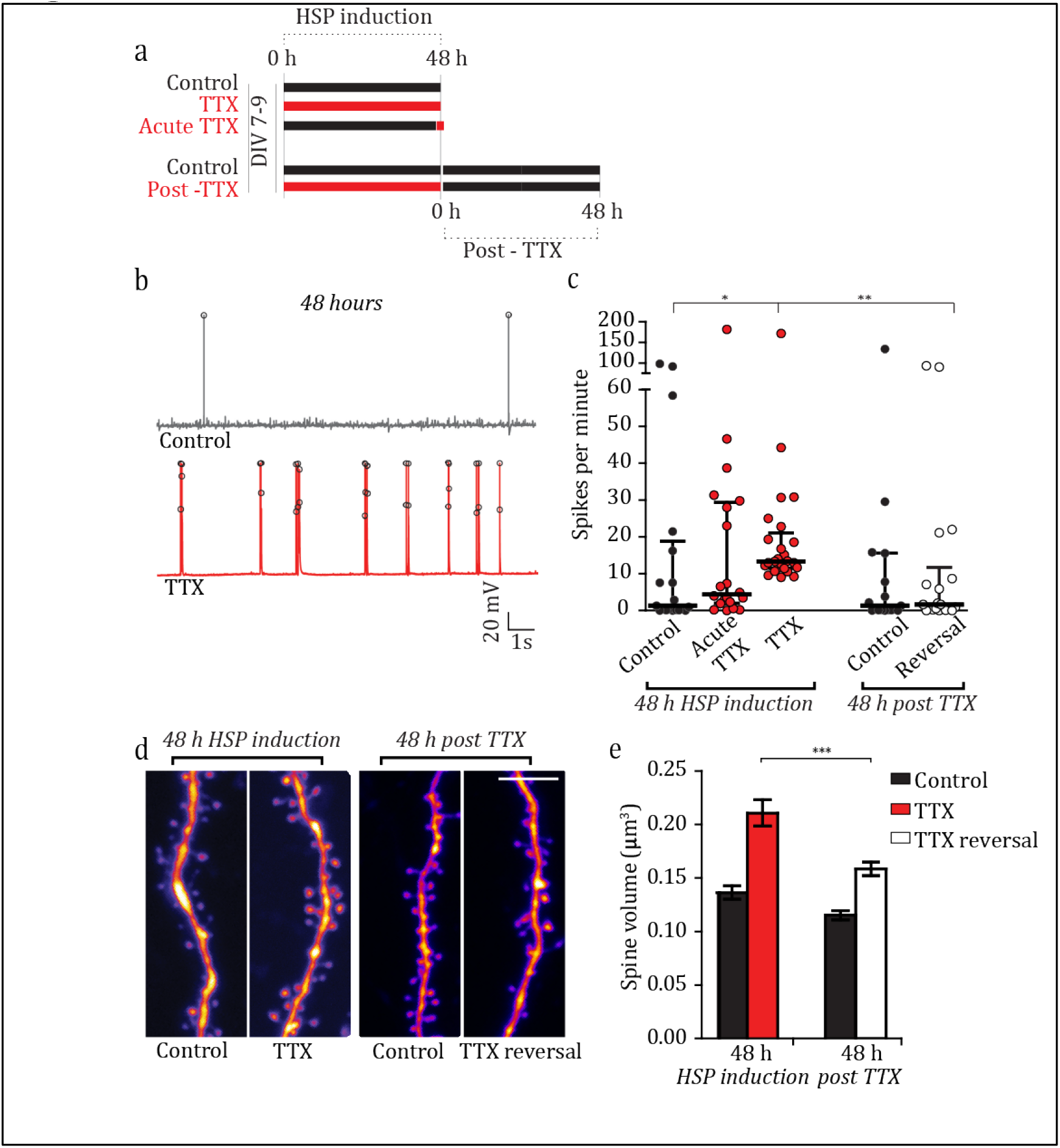
Structural and functional changes after HSP are reversible. Experimental timeline for reversal experiments. b) Example traces showing cell firing following 48 h of activity block. Spikes are demarcated with black circles. c) Quantification of spikes per minute throughout experiment. *p_48 control – 48 TTX_ = 0.03, Kruskal-Wallis post hoc Dunn. **p_48 TTX – 48 post TTX_ = 0.010, Kruskal-Wallis post hoc Dunn. Number of neurons: 48h TTX condition: 48 h Control = 17, 48 h Acute TTX = 20, 48 h TTX = 26; 48h ‘post-TTX’ Control = 15, 48 h post-TTX Reversal = 18. d) Representative images from dendrites after 48 h of HSP induction and at 48 h post TTX. Scale bar = 5μm. e) Quantification of spine sizes throughout the experiment. Graph is plotted as mean ± SEM. All p values calculated using Kruskal-Wallis test, post-hoc Dunn’s. ***p_48 TTX – 48 post TTX_ = 4.1852e-07. Number of spines (n = first number) of Neurons (N = in parentheses): 48 h Control = 285 (7), 48 h TTX = 208 (6), 48 h ‘post-TTX’ Control = 514 (14), 48 h post-TTX = 430 (15).

### Enhanced Hebbian potentiation of small spines after HSP

Functional plasticity can be enhanced upon homeostatic modulation in hippocampal slices (Arendt et al., 2013; Félix-Oliveira et al., 2014), but occlusion of LTP may also occur (Soares et al., 2017). Given our finding that robust growth of spines occurs during homeostatic plasticity, we wanted to determine whether individual inputs were able to undergo further activity-dependent structural plasticity. We therefore induced HSP in hippocampal slices for 48 h and followed this with synaptic potentiation at visually identified dendritic spines through 2-photon mediated glutamate uncaging (Figure 3a,b). This paradigm elicits potentiation and growth of only the stimulated input (Govindarajan et al., 2011; Harvey and Svoboda, 2007). We followed the growth of spines relative to their average baseline volumes, in both the TTX and control conditions (Figure 3c,d). During the initial 45 minutes post-stimulation, both TTX-treated and control spines grew to a similar extent, indicating that homeostatic structural scaling does not occlude activity-dependent structural plasticity (mean ± SEM; Control stimulated 45’= 1.45 ± 0.10; Control neighbors 45’= 1.08 ± 0.04; TTX stimulated 45’= 1.59 ± 0.14; TTX neighbors 45’= 1.14 ± 0.04; p > 0.99) (Figure 3d). However, after 120 minutes, only the stimulated inputs in the TTX-treated neurons remain significantly larger than their neighbors (Figure 3d; mean ± SEM: TTX stimulated 120’= 1.53 ± 0.13; TTX neighbors 120’= 1.16 ± 0.04), as control spines begin to return to their initial volume (Figure 3d; Control stimulated 120’= 1.30 ± 0.01; Control neighbors 120’= 1.14 ± 0.04). Although the TTX and control spines are not significantly different at 120’ (p = 0.1345, Mann Whitney), they begin to diverge, as seen relative to their respective neighbors. Thus, at scaled synapses, activity that would otherwise lead to short lived structural plasticity now elicits long lasting structural plasticity.

**Figure 3.**
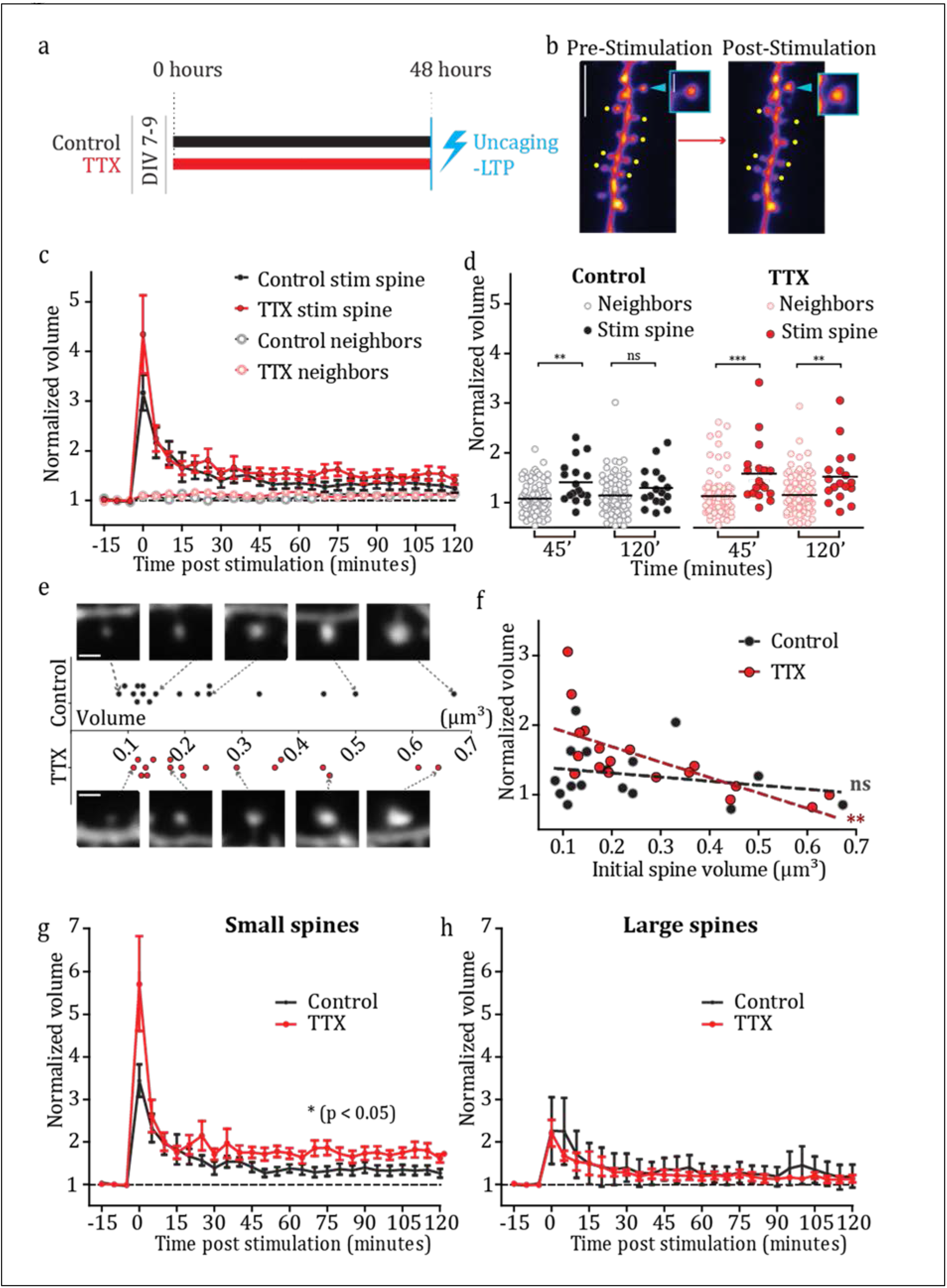
Potentiation of single inputs is enhanced at smaller spines following HSP. Timeline for uncaging experiments. b) Representative images of structural plasticity following single spine glutamate uncaging. Arrowheads and dots indicate stimulated and neighboring non-stimulated spines respectively. Scale bar = 5μm (main image), 1 μm (inset). c) Normalized volume of stimulated and neighboring spines for control and TTX-treated conditions over 2 hours of LTP induction. Volumes were calculated from Z-stacks taken in 5 minute bins from time point 0 (thus the stack represented as 0 was collected between 0-5 minutes, for example). For all data, the total number of spines are represented by the first number (n), and followed by the number of independent neurons (N = in parentheses) from which they were collected. Control Stimulated = 17 (17), TTX Stimulated = 18 (18), Control neighbors = 85 (17), TTX neighbors = 102 (18). d) All analyzed stimulated and neighboring spine volumes at time points 45 minutes and 120 minutes post stimulation. Significance was calculated using one-way ANOVA post-hoc Bonferroni. Control condition: **p_stimulated45 – neighbors45_ = 8.4246e-04. p_stimulated120 – neighbors120_ = 0.6881. TTX condition: ***p_stimulated45 – neighbors45_ = 2.9383e-04. **p_stimulated120 – neighbors120_ = 0.0046. (n same as in panel c). e) Initial absolute spine size vs. normalized spine volume at 120 minutes. TTX spines linear regression: r^2^ = 0.48, **p = 0.002. Control linear regression: r^2^ = 0.06, p = 0.346. f) Distribution of all initial volumes of stimulated spines, with example spine images. The cut-off for defining ‘large’ spines was the median volume off all spines multiplied by 1.5. Scale bar = 1 μm. g) Time-course of the structural LTP experiment plotting only the small spines. *p = 0.0458, 2-way repeated measures ANOVA). Number of spines (n) from neurons (N = in parentheses): Control = 13 (13), TTX = 11 (11). h) Time-course of structural LTP plotting only large spines. p = 0.699, 2-way repeated measures ANOVA. Number of spines (n) from neurons (N = in parentheses): Control = 4 (4), TTX = 17 (17).

Structural plasticity at individual inputs varies with activity and spine size. While weak stimulations preferentially affect small spines (Matsuzaki et al., 2004), strong stimuli can elicit structural plasticity across a variety of spine sizes (Govindarajan et al., 2011; Oh et al., 2013; Ramiro-Cortés and Israely, 2013). Our stimulated spines spanned a range of sizes, allowing us to test whether size interacted with prior expression of HSP when expressing synaptic potentiation and further spine growth (Figure 3e). We examined the amount of structural plasticity expressed by spines of different initial sizes after activity blockade. We found a negative correlation between a spine’s initial average value and its normalized final volume when the spine population had first undergone homeostatic plasticity, but not in the control population, indicating that spine volume is an important modulator of the potential for plasticity after HSP (Figure 3ef; r^2^ = 0.48 for TTX, p < 0.01, r^2^ = 0.06 for control, p > 0.3). To further examine how HSP modulates the size dependence of structural plasticity, we classified spines as ‘large’ if their initial volume was more than 150% of the median initial volume of all spines, and the remainder were placed in the ‘small’ category. Among the small spines, we observed a significant increase in the magnitude of LTP in the TTX condition compared to the controls (Figure 3g; p=0.046, 2-way repeated measures ANOVA). On the other hand, large spines expressed a short-lasting growth that quickly decayed to baseline and was not dependent on whether they had been subjected to activity blockade (Figure 3h; p=0.70 2-way repeated measures ANOVA). Therefore, small spines preferentially underwent long lasting structural plasticity after HSP. We further observed that small spines, which had undergone HSP, tended to show a greater degree of initial growth in response to the induction of plasticity (Figure 3g, time point 0). To quantify the dynamic growth of these spines, we analyzed high-speed images of the spine head taken throughout the stimulation period (approximately 20 Hz) and did not observe a significant difference in the growth curves between the control and TTX treated spines (Figure 4a,b). When we examined the rate of growth that each spine expressed in the first minute after the stimulation however, we found that TTX treated small spines grew more than control small spines (Figure 4c; p = 0.018, 1way ANOVA with post-hoc Tukey’s), while large spines showed no difference between the two conditions. Therefore, glutamate stimulation leads to a sustained growth of homeostatically modified small spines, which may reflect prolonged signaling at these synapses. Together, these data indicate that HSP facilitates Hebbian plasticity, and that this is accomplished at the individual spine level through preferential structural plasticity of small inputs.

**Figure 4.**
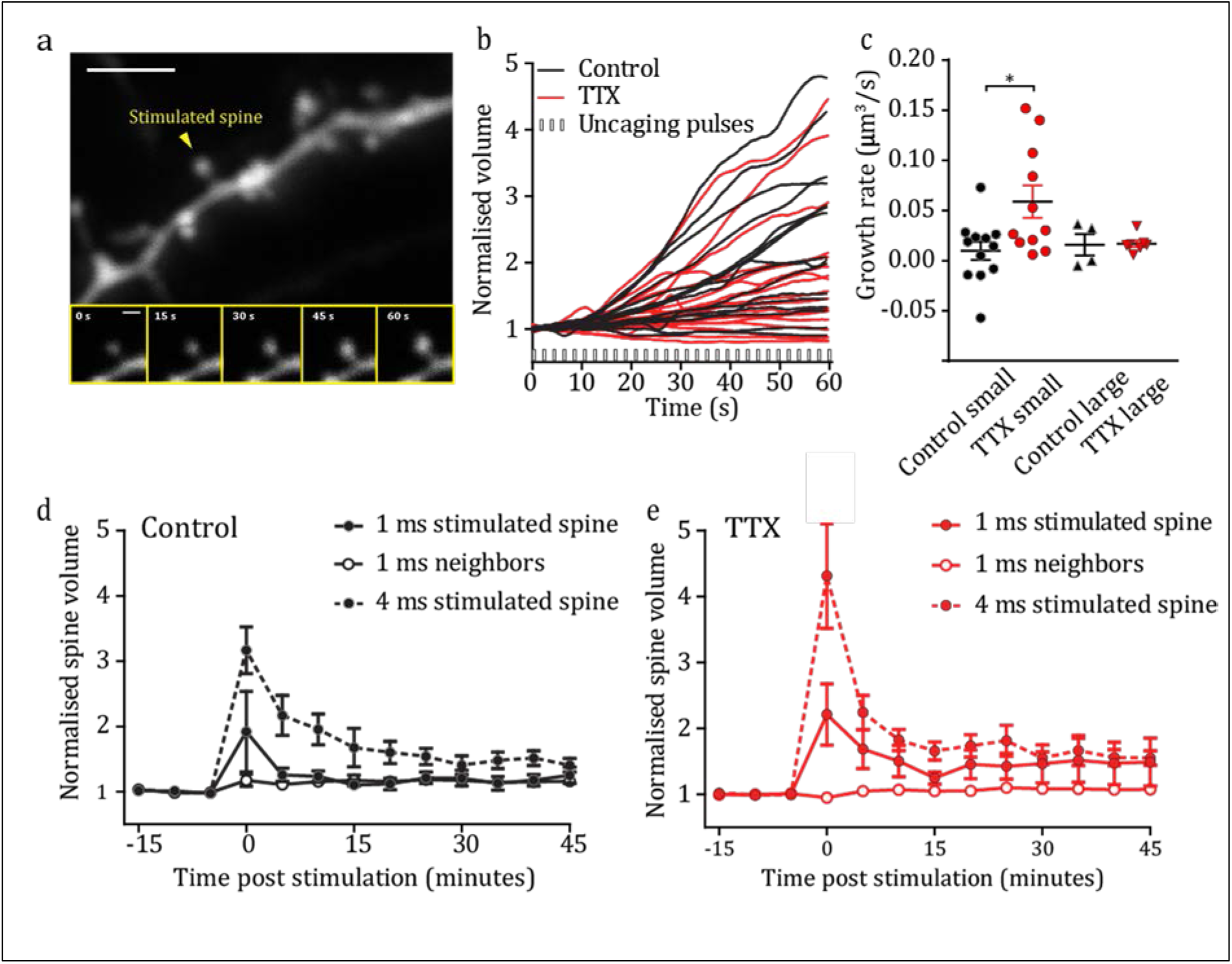
HSP increases the efficacy and reduces the threshold for induction of LTP at individual spines. Representative image of a single spine during the course of uncaging. Scale bar = 5μm (main image), 1μm (inset). b) Individual traces of spine growth during glutamate uncaging stimulation. Control spine growth vs TTX spine growth: p = 0.198, 2-way repeated measures ANOVA. Number of spines (n): Control = 16, TTX = 18. c) Growth rates of stimulated spines for 1 minute after the end of stimulation. *p_small control–small TTX_ = 0.018, 1-way ANOVA with post-hoc Tukey’s. p_large control–large TTX_ < 0.99, 1-way ANOVA with post-hoc Tukey’s. Number of spines (n): Control small = 13, TTX small = 11, Control large = 4, TTX large = 7. d) Quantification of normalized spine volume with different stimulation pulse durations in the control condition. p_1ms stimulated–1ms neighbors_ = 0.292, 2-way repeated measures ANOVA post-hoc Bonferroni. Number of spines (n) from Neurons from which they were collected (N = in parentheses): stimulated 1ms = 11 (11), neighbors 1ms = 52 (11), stimulated 4ms = 17 (17). *_p1ms stimulated–4ms stimulated_ = 0.039 2-way rm ANOVA. Note 4ms data is the same as shown in Figure 3. e) Quantification of normalized spine volume with different stimulation pulse durations in the TTX condition. **p_1ms stimulated–1ms neighbors_ = 0.007, 2-way rm ANOVA. p_1ms stimulated–4ms stimulated_ = 0.27, 2-way rm ANOVA). Number of spines (n) from Neurons (N): stimulated 1ms = 11 (11), neighbors 1ms = 52 (11), stimulated 4ms = 18 (18). Note 4ms data is the same as in Figure 3.

### Homeostatic plasticity facilitates the induction of Hebbian structural plasticity

Having observed that small spines showed enhanced responses to stimulation after homeostatic plasticity, as reflected by a higher rate of growth and longer lasting structural plasticity (Figure 3g and 4c), we reasoned that these inputs may respond more robustly to a weaker stimulation. This is also supported by previous findings that loss of activity at individual inputs lowers their threshold for subsequent plasticity (Lee et al., 2010). To test this possibility, we applied a sub-threshold stimulation that utilizes a shorter laser pulse of 1 ms (instead of 4 ms) during the glutamate uncaging protocol, since this does not induce structural plasticity at spines (Govindarajan et al., 2011; Harvey and Svoboda, 2007). Indeed, control spines that underwent this sub-threshold stimulation showed only a transient post-stimulus growth that lasted 5 min, resulting in no long term plasticity (Figure 4d; Control spines, 1 ms stimulated vs neighbors: p > 0.05, 2-way repeated measures ANOVA, post hoc Bonferroni). These results were significantly different from what we had observed in response to the 4ms stimulation (Figure 4d; 1 ms stimulated vs 4 ms stimulated; p = 0.04, 2-way repeated measures ANOVA; compared to data from Figure 3c). In contrast to this, HSP modified spines grew significantly in response to the subthreshold stimulation and remained larger than their neighbors (Figure 4e; TTX spines, 1 ms stimulated vs neighbors p = 0.01 2-way repeated measures ANOVA, post hoc Bonferroni). Following TTX treatment, both weak and strong stimulations elicited similar levels of long lasting growth of spines (Figure 4e; 1ms stimulated vs 4 ms stimulated: p = 0.27, 2-way repeated measures ANOVA). Thus, homeostatic plasticity facilitates structural plasticity by lowering the threshold for stimulation, without affecting the magnitude of plasticity. In this way, HSP acts locally, in conjunction with activity, to increase the likelihood that an individual synapse will participate in the encoding of information.

### HSP influences structural plasticity of neighboring spines

In addition to modifying synaptic inputs through scaling, homeostatic plasticity can also influence cellular firing rates by modulating the intrinsic excitability of neuronal membranes (Desai et al., 1999). We reasoned that such alterations could facilitate the expression of structural plasticity not only at stimulated spines but also at nearby inputs within the dendritic branch. To test this idea, we examined whether homeostatic plasticity altered neighboring spine volume dynamics following the potentiation of single inputs. We ranked neighboring spines according to their distance from the stimulated spine (with 1 representing the closest neighbor to a stimulated spine) and plotted their growth dynamics for two hours following the stimulation (Figure 5a). We found that spines located in close proximity to the stimulated spine increased in volume only in neurons that had first undergone homeostatic plasticity (compare the first 20 spines, from 0 to 60 minutes after stimulation, Figure 5a). We classified the unstimulated neighbors as “near” or “far” - located either within or beyond 5 μm of the stimulated spine respectively - and quantified spine volume changes over time (Figure 5b-d). As expected, none of the neighbors of stimulated spines changed significantly from their original size in untreated neurons (Figure 5b,c). However, after HSP, neighbors that were within 5 μm of the target spine exhibited significant growth in the 5 minutes following the stimulation (Figure 5b) and remained significantly larger than more distant spines (Figure 5b,d; p = 0.001, 2-way repeated measures ANOVA). Interestingly, farther neighbors (located up to 20 μm away from the site of stimulation) tended to decrease in size for several minutes following activity, although this was not significantly different from baseline (Figure 5d). We next investigated whether there was a correlation between the structural dynamics of each stimulated spine and its neighbors. We calculated correlation coefficients between the volume change of the stimulated spine and those of its neighbors across the time-course of the experiment. While fluctuations in the volumes of neighboring spines in control conditions were independent of the stimulated input (Figure 5e; r2 = 0.00263, p = 0.61), spine volumes in TTX treated neurons changed in accordance with the stimulated spine (Figure 5f; r2 = 0.156, p < 0.001). Specifically, near TTX neighbors had a positive correlation with the stimulated input, reflecting growth, which eventually became a negative correlation at further distances from the stimulated spine. These findings indicate that HSP reduces the input specificity of activity in a distance dependent manner that is at once both cooperative with near, and competitive with far, inputs. This allows for homeostatic modulation to effect structural plasticity within a dendritic region through activity at a single spine, which favors clustered plasticity of synaptic inputs.

**Figure 5.**
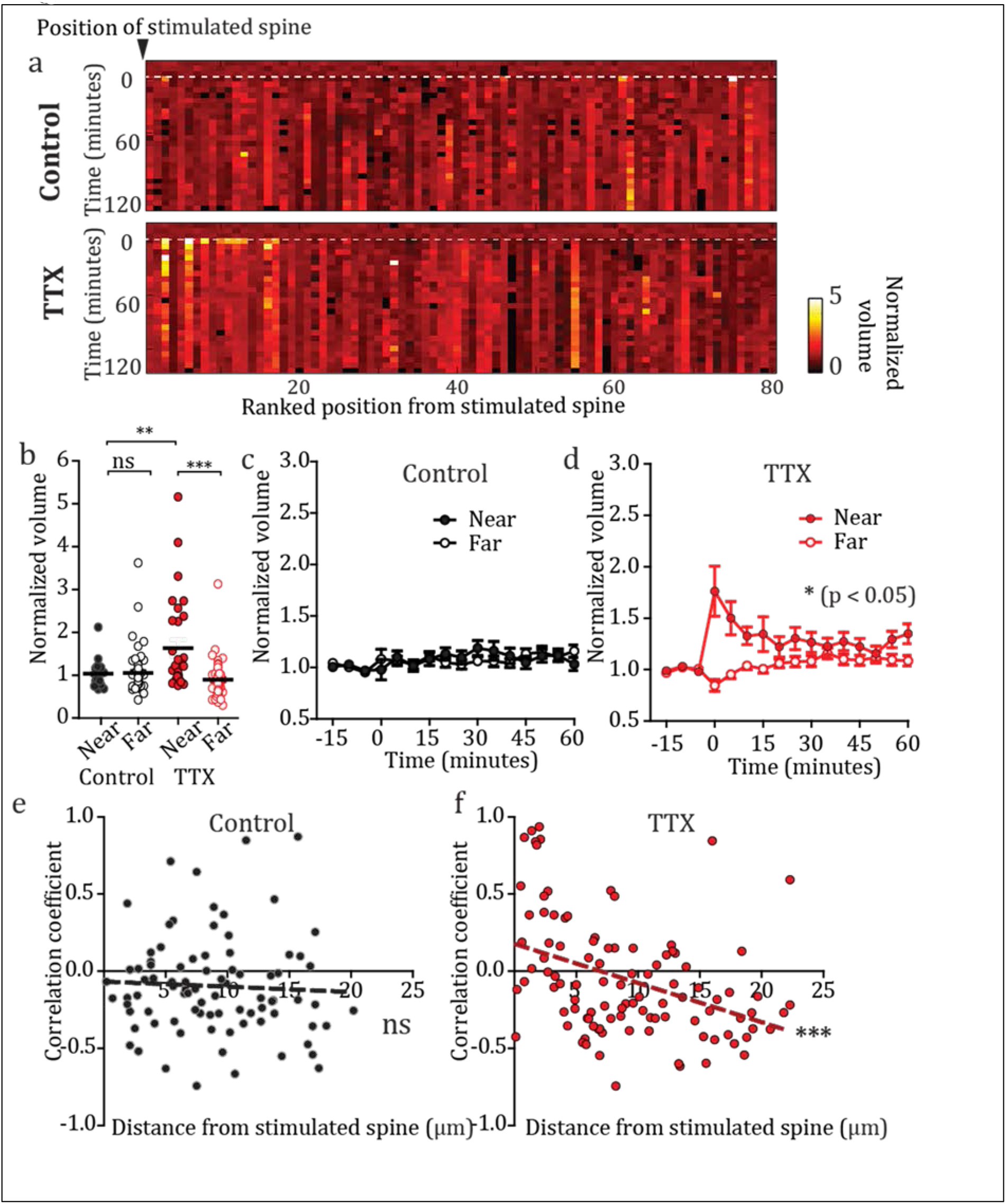
Stimulation of single inputs leads to plasticity of clustered neighbors after HSP. a) Heat maps of growth dynamics of neighboring spines over time. Neighbors from all experiments were pooled and ranked in order distance from the stimulated spine. Index 1 is the spine nearest to its stimulated spine, and 80 is the furthest away. Heat map = normalized spine volume. Number of spines (n) from Neurons from which they were collected (N = in parentheses): Control = 80 (17), TTX = 80 (18). b) Mean volumes for the first 10 minutes following stimulation for the different neighboring conditions. ***p_TTX near – TTX far_ = 1.0674e-05, Kruskal-Wallis. Number of spines (n) from Neurons from which they were collected (N = in parentheses): Control near = 22 (17), control far = 63 (17), TTX near = 32 (18), TTX far = 70 (18). c) Normalized volumes of near and far neighboring spines in the control condition. p = 0.987, 2-way rm ANOVA. Number of spines – same as above. d) Normalized volumes of neighboring spines in the TTX condition. *p = 0.0124, 2-way repeated measures ANOVA. Number of spines – same as above. e) & f) Correlation coefficients of volume change of stimulated spines vs volume change of neighboring spine, plotted against the distance between the neighboring spine and the stimulated spine. e) Linear regression control: r^2^ = 0.003, p = 0.641. f) Linear regression TTX: r^2^ = 0.156, ***p = 3.91e-05).

## Discussion

Synaptic networks must balance the need to enhance efficacy during learning without saturating their capacity for further changes. Homeostatic plasticity can achieve this regulatory role by effecting a gain modulation on neurons in order to maintain synaptic activity within an optimal target range. It is unclear what are the consequences of implementing HSP for the network, and specifically, how its interaction with Hebbian plasticity impacts the subsequent encoding of information at the level of single inputs. Due to the linear relationship between synaptic structure and function, we used live 2-photon imaging and glutamate uncaging to probe the physical consequences of inducing homeostatic plasticity on a neuron, and to determine how this impacts the ability of single synapses to undergo further changes in efficacy. We show that homeostatic synaptic plasticity, induced by 48 hours of activity blockade, leads to the reversible growth of dendritic spines. We demonstrate that this form of modulation facilitates subsequent structural plasticity at single synapses and that this effect preferentially occurs at smaller inputs. After HSP, activity elicits a faster growth rate at these spines, and converts an otherwise subthreshold stimulation into one that is now capable of eliciting structural plasticity. Interestingly, we find that the induction of Hebbian plasticity on a background of homeostatic plasticity leads to compromised input specificity, as neighboring spines grow when they are located in close proximity to a stimulated synapse. Taken together, our results show that homeostatic plasticity can modulate a neuron’s response to activity by facilitating the sensitivity of smaller inputs and inducing structural plasticity at neighboring synapses.

The structural scaling that we observe after 48 hours of activity blockade results in a non-linear upscaling of spines and an increased number of large spines (Figure 1g). By 72 hours, this structural scaling becomes linear (Figure 1k), suggesting that the initial changes may represent a physical overshooting of the target size. This has also been observed on short timescales following glutamate stimulation of single spines, where an initial large volume change is followed by stabilization of the spine at a more modest size (Matsuzaki et al., 2004). Upon the reinstatement of activity, we find that spines return to their original size as firing patterns normalize by 48 hours (Figure 2c-e), indicating that homeostatic structural changes are plastic and reversible, similarly to homeostatic responses to fluctuating activity levels.

We demonstrate that activity blockade increases spine sizes across the population, raising the question of whether further enhancement of potentiation and spine growth could be achieved across inputs of different sizes. Both enhancement and occlusion of plasticity have been reported following HSP at the population level (Arendt et al., 2013; Félix-Oliveira et al., 2014; Soares et al., 2017), but it is unclear how individual inputs are modulated. Further, while larger synapses have been shown to undergo functional homeostatic modulation (Thiagarajan et al., 2005), inputs show a size dependent variation in their ability to undergo Hebbian structural plasticity, large spines being more stable and requiring stronger stimulation in order to be potentiated (Govindarajan et al., 2011; Matsuzaki et al., 2004). We probed plasticity with glutamate uncaging and found that HSP leads to structural plasticity lasting over two hours preferentially at small spines, while control spines decay to baseline after one hour (Figure 3g,h). This suggests that a long lasting, protein synthesis dependent form of Hebbian plasticity was induced at these inputs (Govindarajan et al., 2011), supported by recent findings that TTX-mediated activity blockade leads to the specific production of plasticity related proteins (Schanzenbacher et al., 2018). Closer examination of the structural plasticity we induced revealed that the population of small spines express the majority of the growth, while large spines remain stable (Figure 3g,h). This result implies that plasticity at large spines is saturated after homeostatic modifications, and that their threshold for structural plasticity is not altered by HSP.

Upon stimulation, we did not observe a significant difference in the magnitude of the potentiation that spines expressed, nor in their immediate response to the stimulation itself, but rather we found that spines post-HSP express a significantly faster growth rate compared to their counterparts in the first minute after stimulation (Figure 4a-c). This enhanced response may be due to amplified signaling cascades shared between these forms of plasticity (Fernandes and Carvalho, 2016), which could facilitate the induction of long lasting structural plasticity. It was therefore not surprising to find that a subthreshold stimulation elicits long lasting growth only at inputs that have undergone homeostatic plasticity (Figure 4 d,e), without affecting the initial magnitude of the response between the HSP modified synapses and the control populations. Changes to NMDA receptor composition that occur following activity blockade could provide a means by which to enhance the calcium permeability of synapses and thus would allow for prolonged signaling responses (Barria and Malinow, 2005; Lee et al., 2010; Paoletti et al., 2013). Taken together, these data show that the effects of homeostatic plasticity preferentially impact small spines during Hebbian regulation of synaptic strength by predisposing them to undergo long lasting structural changes. Such modulation may result in the conversion of weaker stimuli into more salient forms, and proposes a more global role for HSP in information encoding beyond the optimization of neuronal activity.

A hallmark of activity dependent changes during Hebbian plasticity is input specificity (Barrionuevo and Brown, 1983). In contrast, homeostatic plasticity is generally expressed over a wider synaptic range, and how this modulation impacts Hebbian learning rules at individual inputs is unknown (Vitureira and Goda, 2013). Computational studies have postulated that homeostatic plasticity can influence previous Hebbian events at a synapse (Rabinowitch and Segev, 2008). Additionally, several lines of evidence indicate that activity can lead to local homeostatic changes within dendrites (Branco et al., 2008; Ju et al., 2004; Liu, 2004; Sutton et al., 2006). We therefore considered the possibility that HSP may function to influence subsequent Hebbian plasticity. Indeed, we find that HSP induces the spillover of Hebbian structural plasticity within the dendrite, as seen by the growth of neighboring unstimulated spines following the activation of one input (Figure 5). Thus, homeostatic plasticity drives the local expression of Hebbian plasticity and reduces input specificity. While cooperative interactions have been shown to occur between inputs when multiple synapses are stimulated (Govindarajan et al., 2011; Harvey and Svoboda, 2007), in the absence of such coactivation, structural changes at neighbors have thus far served to counterbalance the direction of plasticity (Oh et al., 2015). Our observations that HSP promotes activity-driven growth of a group of synapses within a five micron area is in alignment with the observed distance over which plasticity related proteins can spread between co-active synapses (Harvey et al., 2008). Thus, the interaction between homeostatic and Hebbian plasticity may coordinately delimit a region of clustered plasticity, which has been proposed to increase the memory capacity of a neural circuit (Chklovskii et al., 2004; Govindarajan et al., 2011, 2006; Ramiro-Cortés et al., 2014). Although clustered activity and its structural correlates have begun to emerge (Fu et al., 2012; Kleindienst et al., 2011; Makino and Malinow, 2011; Yadav et al., 2012), the mechanisms which drive the physical organization of inputs are unknown. HSP may provide a basis for such organization by reducing the threshold for cooperativity between inputs, allowing primed synapses located within several microns of an active spine to be more easily potentiated. This may serve as a first step in the physical arrangement of synapses into clusters. The broader consequence of this could be the binding together of information of varying saliencies, within a delimited region, into the same engram.

Our finding that synaptic threshold modulation after HSP is implemented at small, rather than at large spines, and in a reversible manner, suggests that synaptic scaling mechanisms are separable from plasticity mechanisms. However, by promoting long-lasting plasticity at select spines, and potentiating unstimulated neighboring spines within a delimited dendritic region, homeostatic plasticity may interact with and reduce the input specific nature of Hebbian plasticity, enhance the clustering of synaptic inputs, and shape the long-term organization of neural circuits. In this way, despite the global nature of homeostatic plasticity, inputs may be locally modulated in an activity-dependent manner.

## Materials and Methods

### 1. Slice culture preparation and biolistic transfection

Mouse hippocampal organotypic slice cultures were prepared using p7-10 C57BL/6J mice as previously described (Govindarajan et al., 2011). Briefly, hippocampi were dissected and 350 μm slices were cut with a tissue chopper in ice-cold artificial cerebral spinal fluid (aCSF) containing 2.5 mM KCl, 26 mM NaHCO_3_, 1.15 mM NaH_2_PO_4_, 11 mM D-glucose, 24 mM sucrose, 1 mM CaCl_2_ and 5 mM MgCl_2_. The slices were cultured on membranes (Millipore), and maintained at an interface with the following media: 1× MEM (Invitrogen), 20% horse serum (Invitrogen), 1 mM GlutaMAX (Invitrogen), 27 mM D-glucose, 30 mM HEPES, 6 mM NaHCO_3_, 2mM CaCl_2_, 2mM MgSO_4_, 1.2% ascorbic acid, 1 μg/ml insulin. The pH was adjusted to 7.3, and the osmolarity adjusted to 300–310 mOsm. All chemicals were from Sigma unless otherwise indicated. Media was changed every 2-3 days.

Biolistic transfection of slice cultures was accomplished using a Helios gene gun (Bio-Rad) after 4–7 days in vitro (DIV). Gold beads (10 mg, 1.6 μm diameter, Bio-Rad) were coated with 100 mg AFP-plasmid DNA (a GFP-expressing plasmid driven by the β-actin promoter (Inouye et al., 1997) according to the manufacturer’s protocol and delivered biolistically to the slices, using a pressure of 160-180 psi.

### 2. Homeostatic plasticity induction by activity block

TTX (1 μM) was added to the culture media at 7-9 DIV. The day of application was then designated day 0 for experiments. Control experiments were maintained in normal culture media, and were age- and animal-matched to treated slices for experiments.

### 3. Patch Clamp Electrophysiology

Hippocampal slice cultures were perfused continuously with aCSF (as above, with the addition of 0.5 μM TTX for all mEPSC recordings) for a pre-incubation period of 15 to 30 min. Whole cell voltage-clamp recordings were performed in CA1 pyramidal neurons, using 7–8 ΜΩ electrodes. For mEPSC recordings, the internal solution contained 135 mM Cs-methanesulfonate, 10 mM CsCl, 10 mM HEPES, 5 mM EGTA, 2 mM MgCl_2_, 4 mM Na-ATP and 0.1 mM Na-GTP, with the pH adjusted to 7.2 with KOH, at 290-295 mOsm. Cells were voltage clamped at −65 mV. Cellular recordings in which series resistance was higher than 25 mV were discarded. Stability was assessed throughout the experiment, with cells whose series resistance changed more than 30% being discarded. mEPSCs recordings were started 3 minutes after break-in and continued for 10 minutes. Signals were acquired using a Multiclamp 700B amplifier (Molecular Devices), and data was digitized with a Digidata 1440 at 3 kHz. mEPSC events were detected off-line using Mini-Analysis Program (Synaptosoft). Events smaller than 15 pA fell within the range of noise in the system and were not included in the analysis. For spontaneous activity recordings, slices were perfused continuously with aCSF without the addition of TTX for a pre-incubation period of 5 to 10 min. The internal solution for the electrodes contained 136.5 mM K-Gluconate, 9 mM NaCl, 17.5 mM KCl, 10 mM HEPES, 0.2 mM EGTA, and 0.025 mM Alexa 594, with the pH adjusted to 7.2 with KOH, at 280-290 mOsm. In current clamp with no external current applied, an IV curve was first recorded to check for spike frequency accommodation in order to validate the identity of pyramidal neurons. Spiking events were then recorded for a period of 6-9 minutes. The addition of Alexa-594 allowed cells to be imaged post-recording.

### 4. Two-photon Imaging

Two-photon imaging was performed on a BX61WI Olympus microscope, using a galvanometer-based scanning system (Prairie Technologies /Bruker) with a Ti:sapphire laser (910 nm for imaging AFP; Coherent), controlled by PrairieView software (Prairie Technologies). Slices were perfused with oxygenated aCSF containing 127 mM NaCl, 2.5 mM KCl, 25 mM NaHCO_3_, 1.25 mM NaH_2_PO_4_, 25 mM D-glucose, 2 mM CaCl_2_, 1 mM MgCl_2_ and 0.5 μM TTX (equilibrated with O_2_ 95%/CO_2_ 5%) at room temperature, at a rate of 1.5 ml/min. Secondary or tertiary apical dendrites of CA1 neurons (where the apical trunk is counted as the primary branch) were imaged using a water immersion objective (60×, 1.0 NA, Olympus LUMPlan FLN) with a digital zoom of 8×. For each neuron, 2-3 dendrites were imaged. Z-stacks (0.5 μm per section) were collected at a resolution of 1024 × 1024 pixels, resulting in a field of view of 25.35 × 25.35 μm. 3 images were taken per dendrite at 5-minute intervals, with the reported spine volume being the average of the 3 images. Images were taken at the highest possible fluorescence intensity without image saturation in order to accurately quantify spine volumes (see below).

### 5. Glutamate uncaging

Uncaging experiments (Pettit et al., 1997) and caged glutamate calibration were carried out as previously described (Govindarajan et al., 2011), and briefly as follows. MNI-caged-L-glutamate (MNI-Glu) (Tocris) was reconstituted in the dark in aCSF lacking MgCl_2_ or CaCl_2_ to make a 10 mM stock solution. Individual aliquots were diluted to the working concentration of 2.5 mM MNI-Glu in uncaging aCSF (see below), in 3 ml volumes. We tested each batch of reconstituted MNI-Glu as previously described (Govindarajan et al., 2011). Briefly, five uncaging test pulses of 1 ms were delivered to single spines and evoked EPSCs were measured by whole cell patch clamp recordings. We compared these to spontaneous mEPSCs, and determined the power needed (60mW at the back aperture) to produce an uncaging EPSC of comparable size to that of an average spontaneous mEPSC. During plasticity experiments, slices were incubated for 30-45 minutes in normal aCSF as described above. Uncaging was performed using a Ti:sapphire laser (720 nm; Coherent), controlled by PrairieView software (Prairie Technologies, Bruker). All stimuli were carried out in uncaging-aCSF containing: 2.5 mM MNI-Glu, 0 mM MgCl_2_, and 4 mM CaCl_2_. The uncaging train consisted of 30 pulses at 0.5 Hz, with a pulse width of either 4 ms (standard protocol) or 1 ms (sub-threshold protocol), using 30mW power as measured at the back aperture (half the power as determined in the calibration step, as previously described for an uncaging LTP protocol (Govindarajan et al., 2011)). The uncaging point was positioned 0.6 μm from the end of the spine head, away from the parent dendrite. Each experiment began with a sham stimulation in uncaging-aCSF lacking MNI-Glu on a control dendrite of the experimental neuron, to rule out the possibility of nonspecific structural changes due to phototoxicity or poor neuronal health. Subsequently, experimental spines on new dendrites were identified, uncaging-aCSF containing 2.5mM MNI-glutamate was recirculated for 5 minutes, and the uncaging protocol was delivered. Fast imaging (approximately 20 Hz) of a region of interest (ROI) of the stimulated spine head and neck was performed throughout the 60 second stimulation, to record the growth dynamics for this time period. After the stimulation, the perfusion was returned to normal aCSF for the remainder of the experiment (1-2 hours). The first image was taken immediately after returning to normal aCSF and is designated as time = 0 post stimulation.

### 6. Spine Volume Determination

To quantify spine volume, we used the custom built Matlab plug-in SpineS, which uses semiautomatic detection, automatic alignment and segmentation of spine heads (Erdil et al., 2012; Ghani et al., 2017). Spine volume was calculated using the summed fluorescence intensity of the spine, normalized to the median fluorescence intensity of the parent dendrite, in order to correct for changes in overall fluorescence levels within a cell. Fluorescence intensity was converted to real volumes by taking Full Width Half Max (FWHM) measurements of spines and calculating a conversion factor between the two measures (Matsuzaki et al., 2004). For the homeostatic plasticity analysis (Figures 1 & 2), all spines with a discernible head within the field of view were included in the analysis, unless obstructed by other structures. For the uncaging analysis (Figures 3, 4 & 5), between 2 and 8 neighboring spines were analyzed from the same dendrite, within the whole field of view. Normalization of spine volumes throughout the experiment was performed using the mean of three baseline images.

### 7. Statistical analyses

All statistical analyses were carried out in Graphpad Prism. Stars (*) represent degrees of significance as follows: (*) = p < 0.05; (**) = p < 0.01; (***) = p < 0.001).

## Acknowledgements

We thank Ali Ozgur Argunsah for his advice and help with data analysis. We thank Steven Kushner, Rui Costa, and members of the Israely lab for their support and critical reading of the manuscript. This work was supported by grants from the Bial Foundation (to II), Fundação para a Ciência e a Technologia (to II and AH), and CONACYT 254878 (to YRC).

## Author Contributions

AFH, YRC and II designed the experiments. AFH performed the experiments and analyzed the data. AFH, YRC and II wrote the manuscript.

